# Targeting of CIP4-Calcineurin Signalosomes Improves Cardiac Structure and Function After Myocardial Infarction

**DOI:** 10.1101/2025.05.12.653597

**Authors:** Anne-Maj Samuelsson, Abraham L. Bayer, Jinliang Li, Yang Li, DeEsmond Lewis, Moriah Gildart Turcotte, Kimberly L. Dodge-Kafka, Pilar Alcaide, Michael S. Kapiloff

## Abstract

**Background:** Calcineurin in a pleiotropic signaling enzyme that promotes pathological cardiac remodeling but also cardioprotection in ischemia-reperfusion injury. In addition, calcineurin inhibitors are immunosuppressants. This pleiotropy has precluded the use of calcineurin inhibitors as treatments for heart failure. Cdc42-interacting protein 4 (CIP4/TRIP10) is an endosomal scaffold protein that organizes a calcium and calcineurin Aβ2 (CaNAβ2) signaling compartment activated by G-protein coupled receptors independently of contractile calcium. CIP4 binds CaNAβ2 via the CaNAβ-specific N-terminal polyproline (PP) domain. We previously showed that targeting of CIP4-CaNAβ2 signalosomes inhibited pathological cardiac hypertrophy and the development of heart failure induced by chronic pressure overload in mice. It is unknown whether CIP4-CaNAβ2 signalosomes contribute to cardioprotection and/or cardiac remodeling in ischemic heart disease.

**Methods:** CIP4 conditional knock-out (CKO) mice were studied by echocardiography with strain analysis and histology following ischemia-reperfusion (I/R) injury and permanent left coronary artery (LCA) ligation to induce myocardial infarction. Wildtype C57BL/6NJ mice were transduced with adeno-associated virus (AAV) engineered for cardiomyocyte-specific expression of either a CaNAβ2 shRNA to inhibit CaNAβ2 expression, a VIVIT peptide to inhibit CaN-NFAT signaling, or a CaNAβ2 PP peptide to block CIP4-CaNAβ2 binding. AAV-transduced mice were studied by I/R injury. Additional mice were subjected to permanent LCA ligation and subsequently treated with AAV to test the effects of CaN inhibition in chronic ischemic cardiomyopathy. The effects of CaNAβ2 PP-GFP expression on primary T-cell activation were studied *in vitro*.

**Results:** CIP4 CKO mice and mice expressing the PP anchoring disruptor peptide exhibited preserved cardiac function after I/R injury and decreased infarct size and preserved cardiac function 8 weeks after myocardial infarction by permanent LCA ligation. In contrast, cardiomyocyte-specific depletion of CaNAβ2 and VIVIT peptide expression worsened outcome after I/R injury and in chronic ischemic cardiomyopathy. In addition, in contrast to cardiomyocytes, PP-mediated CaNAβ anchoring inhibition had no effect on T-cell activation and cytokine expression *in vitro*.

**Conclusions:** CIP4-CaNAβ2 signalosomes promote adverse cardiac remodeling and are not cardioprotective. Proof-of-concept is provided for the treatment of ischemic cardiomyopathy by a PP anchoring disruptor gene therapy. Targeting these complexes may be beneficial in cardiovascular diseases, including ischemic cardiomyopathy and acute myocardial infarction.

**Clinical Perspective:** *What is New?:*

- Targeting CIP4, which is a scaffold protein for the phosphatase calcineurin, improves cardiac function in mice after acute myocardial infarction due to ischemia-reperfusion injury and in chronic ischemic cardiomyopathy.
- Gene therapy-based expression of a calcineurin Aβ-derived polyproline peptide, which can compete CIP4-calcineurin binding, is beneficial in acute and chronic myocardial infarction.

*What Are the Clinical Implications?:*

- This study establishes CIP4 signalosomes as a new drug target for the treatment of ischemia-reperfusion injury and chronic pathological cardiac remodeling.
- This study provides proof-of-concept for a new gene therapy approach to treating acute myocardial infarction and chronic ischemic cardiomyopathy.

## Introduction

Heart failure, the common end-stage for cardiac disease, is a syndrome of major public health significance. The prevalence of heart failure among adult Americans is ∼6.7 million, and 1 in 8 will die while in heart failure.^1^ Notably, coronary heart disease is the most common risk factor for heart failure. While current guideline-directed medical therapy for heart failure with reduced ejection fraction (HFrEF) can significantly lower the risk of hospitalization and death,^2^ mortality remains high for all etiologies of heart failure, compelling the development of novel therapeutic strategies.^1^ Pathological cardiac remodeling underlies the development of heart failure. While cardiomyocyte hypertrophy is the major intrinsic compensatory mechanism for chronic stress on the heart, in pathological conditions hypertrophy is accompanied by altered myocyte contractility and metabolism, the progressive loss of myocytes, myocardial inflammation, and interstitial fibrosis, together resulting in systolic and/or diastolic cardiac dysfunction.^3^ This remodeling is regulated by a network of intracellular myocyte signaling pathways that represent potential candidate targets for drug development.^4^

Diverse signaling enzymes have been identified as critical mediators of pathological cardiac remodeling.^4^ One enzyme that is of longstanding interest is the Ca^2+^/calmodulin-dependent phospho-serine/threonine phosphatase calcineurin (CaN, PP2B, PPP3C).^5,6^ In mammals, there are three genes (*PPP3C A-C*) encoding the CaN catalytic A-subunit α, β and γ isoforms.^7^ In addition, CaNAβ is expressed as the alternatively-spliced isoforms CaNAβ1 and CaNAβ2, the latter usually comprising the majority of CaNAβ in cells and sharing a C-terminal domain structure similar to CaNAα.^8^ Studies using mice with CaN Aα and Aβ genetic deletion have revealed isoform-specific functions in different organs.^9–14^ In particular, CaNAβ knock-out attenuated the cardiac hypertrophy induced by pressure overload and angiotensin II and isoproterenol infusion.^9^ The role of CaNAβ in cardiomyocyte hypertrophy is primarily through the action of CaNAβ2, while CaNAβ1 overexpression opposes pathological hypertrophy.^15,16^ Although CaNAβ gene targeting results in the inhibition of pathological cardiac hypertrophy, CaNAβ knockout also increases myocardial loss after ischemia-reperfusion (I/R) injury.^17^ In addition, CaN inhibitors are immunosuppressants, and CaNAβ gene targeting negatively affects T-cell development and activation.^18^ The roles for CaN in cardioprotection and the immune system have precluded the development of CaN-targeting therapeutics for heart failure.

Isoform-specific functions in some tissues may be attributable to differences in expression or substrate specificity.^13,19^ However, differences such as these do not adequately explain why CaN isoforms have a non-redundant function in the adult heart, in which Aα and Aβ (mainly Aβ2) are normally present at similar protein levels and in which relevant substrates can be dephosphorylated with similar efficiency by both isoforms.^9,19^ Specificity in enzyme function can also be conferred by binding to multivalent scaffold proteins that localize the signaling enzyme within the cell and recruit relevant upstream activators and downstream effector substrates. Like other phosphatases,^20^ CaN is bound by scaffold proteins and localized to diverse intracellular compartments.^5,6^ We discovered that in contrast to CaNAβ1, which is intracellularly localized via its different C-terminal domain,^7^ CaNAβ2 can be selectively localized by the CaNAβ-specific N-terminal polyproline (PP) domain to a compartment within the myocyte organized by the endosome-associated scaffold protein Cdc42-Interacting Protein 4 (CIP4, TRIP10).^16^ CIP4 contains a N-terminal Fer-CIP4 Homology (FCH)-Bin/Amphiphysin/Rvs (F-BAR) domain that binds membrane phospholipids and cytoskeletal proteins and confers CIP4 homo-dimerization, a HR1 domain that binds active Rho family members (Cdc42, TC10, and TCL), and a C-terminal Src Homology 3 (SH3) domain that binds proteins involved in the control of the actin cytoskeleton and signaling.^21^ The CIP4 SH3 domain binds the CaNAβ PP-domain.^16^

By imaging live ventricular myocytes expressing fluorescent biosensors, we found that CIP4-bound CaNAβ2 was activated by G-protein coupled receptor signaling, including angiotensin II, α- and β-adrenergic receptors.^16^ Remarkably, CIP4-bound CaNAβ2 was not activated by pacing that induces myocyte contraction. Both CIP4 gene targeting and adeno-associated virus (AAV)-mediated CaNAβ PP peptide expression, which competes CIP4-CaNAβ2 binding, inhibited cardiac remodeling and improved cardiac function in response to pressure overload in mice, apparently independently of nuclear factor of activated T-cells (NFAT) transcription factor regulation.^16^ These results suggested that CIP4-CaNAβ2 “signalosomes” might constitute a new target for intervention in pathological cardiac remodeling. As a major impediment to the development of CaN-directed therapeutics are its roles in cardioprotection and immunity, we were interested whether CIP4-CaNAβ2 signalosomes contribute to these functions, or, alternatively, whether CIP4-CaNAβ2 might be safely targeted in ischemic cardiomyopathy and/or lack a role in T-cells, in which case CIP4-CaNAβ2 signalosomes might be acceptable drug targets for the treatment of heart failure. Here we report the effects of CIP4 cardiomyocyte-specific knock-out and CaNAβ PP anchoring disruption in adult mouse models of I/R injury and chronic ischemic cardiomyopathy. Inhibition of CIP4-CaNAβ2 signalosomes is contrasted with inhibition of CaNAβ2 expression across intracellular compartments by RNA interference and with inhibition of CaN function by expression of a VIVIT peptide that competes the allosteric binding to all CaNA isoforms of substrates like NFAT containing a PxIxIT short linear motif.^7,22–25^ In addition, we explore the effects of PP anchoring disruption in T-cell activation *in vitro* to test for potential adverse immunologic effects. Results are presented that CIP4-CaNAβ2 signalosomes comprise a novel compartment contributing to the deleterious effects of cardiomyocyte CaN signaling in ischemic heart disease that may be therapeutically tractable for the treatment and prevention of heart failure.

## Methods

Complete detailed methods are provided in the Online Supplemental Methods. The data that support the findings of this study are available from the corresponding author upon reasonable request.

### Animal Models

All experiments involving animals were approved by the Administrative Panel on Laboratory Animal Care Institutional Animal Care and Use Committee at Stanford University. The conditional “floxed” *CIP4* (*Trip10*) mouse (Jackson Laboratory Strain #035591) is as previously described.^16^ To induce gene knock-out, *CIP4^f/f^*;Tg(Myh6-cre/Esr1*) “CIP4 CKO” mice and control Tg(Myh6-cre/Esr1*) “MCM” and *CIP4^f/f^*mice were fed tamoxifen-laden chow (125 mg tamoxifen/kg chow, Harlan Teklad) for 1 week before study. For AAV studies, C57BL/6NJ mice (Jackson Laboratory Strain #005304) were injected intravenously with 10^12^ viral genomes either 4 weeks before I/R injury or 2 days after permanent coronary artery ligation to induce myocardial infarction (MI). Echocardiography, strain analysis, and histological analyses were performed as described in the online Supplemental Methods (see also Figures S1 and S2).

### Statistics

Statistics were calculated using Graphpad Prism 10. *n* refers to the number of individual mice. All data are expressed as mean ± s.e.m. Repeated symbols are used as follows: * *p* ≤ 0.05; ** *p* ≤ 0.01; *** *p* ≤ 0.001; **** *p* ≤ 0.0001. For two-group comparisons, following Anderson-Darling (A2*) normality testing, two-tailed t test was performed. For comparisons of multiple groups, datasets failing D’Agostino-Pearson omnibus (K2) normality testing were analyzed by Kruskal-Wallis and Dunn’s multiple comparison tests. Normal multiple group datasets were analyzed by one-way ANOVA followed by Tukey’s post-hoc testing (Brown-Forsythe test was negative for such datasets herein). 2-way ANOVA was used for experiments with matched 2-factor design followed by Tukey’s multiple comparisons (multiple treatment groups) or Uncorrected Fisher’s LSD (two treatment groups) tests.

## Results

### Inhibition of post-myocardial infarction remodeling by CIP4 gene targeting

We previously described the CIP4 conditional knock-out (CIP4 CKO) *CIP4^f/f^*;Tg(Myh6-cre/Esr1*) mouse, in which cardiomyocyte-specific deletion of *CIP4* Exons 2 and 3 results from tamoxifen activation of a “MerCreMer” estrogen receptor – cre recombinase fusion protein (MCM) expressed in cardiac myocytes.^16,26^ CIP4 CKO prevented the development of pathological cardiac remodeling and heart failure in response to chronic pressure overload, apparently in association with the inhibition of CaNAβ2 signaling.^16^ To test whether CIP4-CaNAβ has a broader role in the regulation of pathological remodeling, we now considered whether CIP4 CKO would be similarly beneficial in the chronic ischemic cardiomyopathy associated with myocardial infarction (MI) induced by left coronary artery permanent ligation (LCA PL).^27^ 8-10-week-old CIP4 CKO mice were fed tamoxifen-laden chow for 1 week and then subjected 1 week later to LCA PL or sham survival surgery (Figure 1A). Tamoxifen-treated control cohorts included MCM and *CIP4^f/f^* mice to account for effects of MCM expression and *CIP4* gene *loxP* site insertion, respectively. Consistent with our previously published results,^16^ no significant differences were observed by M-mode echocardiography between CIP4 CKO, MCM and *CIP4^f/f^* mice 4 weeks after sham operation (Figure 1B-E), demonstrating that CIP4 gene deletion has little effect on physiological cardiac function. 4 weeks after MI, MCM and *CIP4^f/f^* mice were found to have left ventricular (LV) dilatation with 25% and 29% increased end-diastolic LV diameter, respectively (Figure 1B,C). Systolic function was impaired with LV end-systolic diameter increased 60% and 62% and fractional shortening less in value by 20% and 18%, respectively for the MCM and *CIP4^f/f^* cohorts (Figure 1D,E). Surprisingly, CIP4 CKO mice did not have significantly increased LV end-diastolic diameter and exhibited less systolic dysfunction than the control cohorts, including a lesser increase in end-systolic LV diameter (24%) and a decrease in fractional shortening of 9%. The relatively preserved systolic function post-MI of the CIP4 CKO mice was confirmed by strain analysis using B-mode parasternal long axis (PS-LAX) images (Figure S3, Videos S1-6). Both segmental and global longitudinal strain were improved by CIP4 CKO following MI when compared to the two control cohorts.

**Figure 1.**
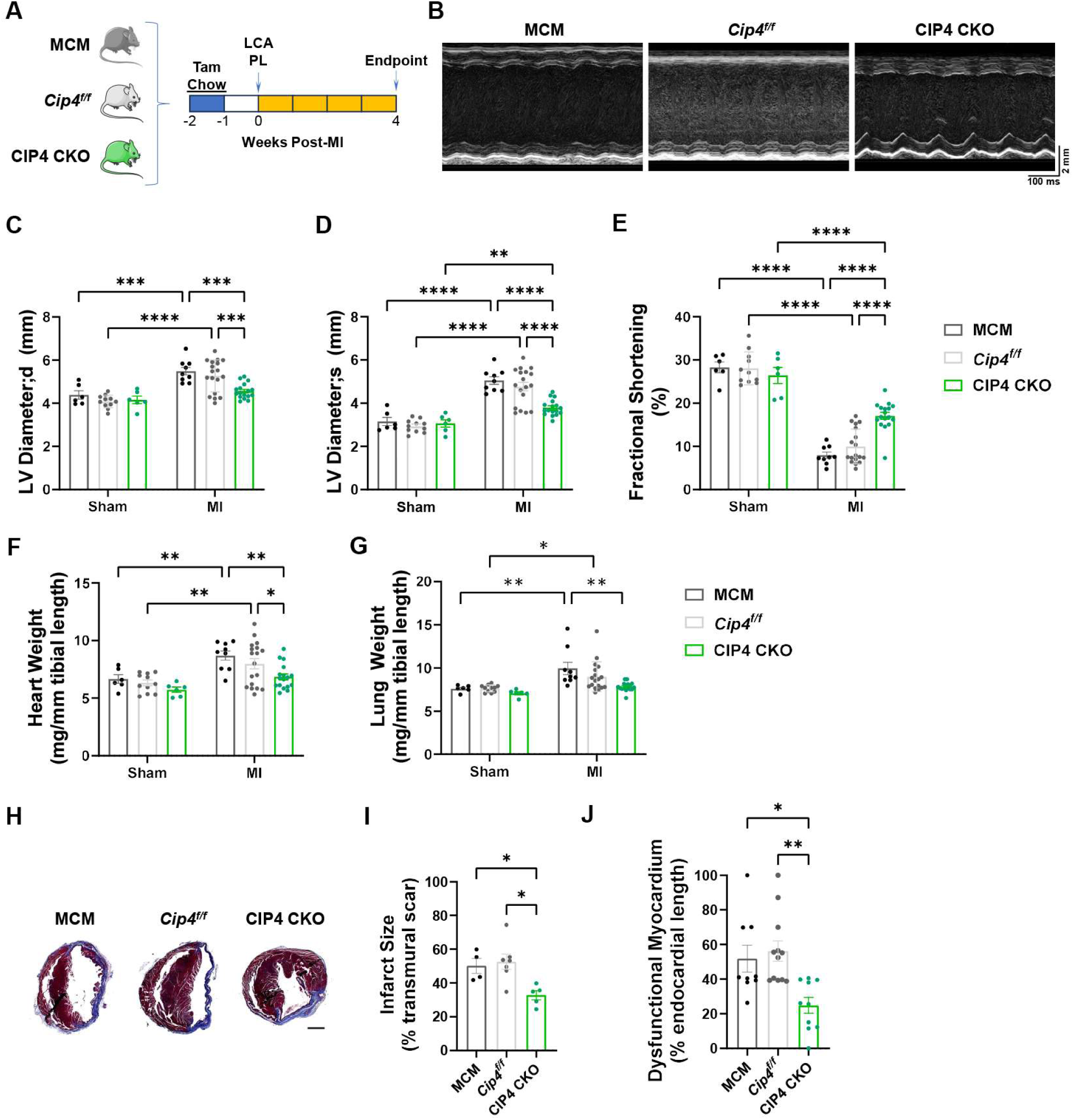
Chronic ischemic cardiomyopathy is improved by *CIP4* conditional gene targeting. **A.** 8-10-week-old male and female CIP4 CKO *CIP4^f/f^*;Tg(Myh6-cre/Esr1*) and control Tg(Myh6-cre/Esr1*) (MCM) and *CIP4^f/f^* mice were treated with tamoxifen-laden chow for one week to induce cre-catalyzed recombination, followed by one week on normal chow before being subjected to permanent ligation of the left coronary artery (LCA PL) or sham surgery. Endpoint studies were performed 4 weeks after induction of myocardial infarction (MI). **B.** Representative M-mode echocardiography at endpoint for MI cohorts. **C-E.** LV diameter in diastole (d) and systole (s) and fractional shortening at endpoint. **F-G,** Gravimetric analysis at endpoint. For C-G, *n*: MCM – Sham – 6; *CIP4^f/f^* - Sham – 11; CIP4 CKO – Sham – 6; MCM – MI – 9; *CIP4^f/f^* - MI – 18; CIP4 CKO – MI – 17. Data in C-F analyzed by 2-way ANOVA and Tukey’s multiple comparison test; data in G analyzed by Kruskal-Wallis and Dunn’s multiple comparison tests. **H.** Representative transverse cardiac sections stained with Masson’s Trichrome. Scale bar – 1 mm. **I.** Infarct size measured as % left ventricular circumference containing >50% transmural scar. *n*: MCM – 4; *CIP4^f/f^* – 7; CIP4 CKO – 5. **J.** Dysfunctional myocardium for MI cohorts is expressed as the ratio of the endocardial length for segments with decreased longitudinal peak systolic strain (see Table S1 for control values) to the overall endocardial length in PS-LAX echocardiographic images (Figures S1). *n*: MCM –9; *CIP4^f/f^* - 12; CIP4 CKO – 10. Data in I and J analyzed by 1-way ANOVA and Tukey’s multiple comparison test. \**p*<0.05; \*\**p*<0.01; *** *p*<0.001; **** *p*<0.0001.

Gravimetric analysis confirmed that CIP4 attenuated post-MI cardiac remodeling. Cardiac hypertrophy (increased biventricular weight indexed to tibial length) was less for the CIP4 CKO mice than MCM and *CIP4^f/f^*cohorts (Figure 1F). In addition, indexed wet lung weight, a marker for heart failure, was increased only for the two control cohorts and not for CIP4 CKO mice (Figure 1G). Notably, the histological measurement of infarct size at endpoint showed that CIP4 CKO reduced infarct size by 35-38% compared to control mice (Figure 1H,I). This result was corroborated by the measurement of fractional dysfunctional myocardium by segmental longitudinal strain analysis (Figure 1J). Together, these results showed that CIP4 CKO preserved cardiac structure and function in a model of chronic ischemic cardiomyopathy, suggesting that like in chronic pressure overload^16^ CIP4 targeting inhibits pathological cardiac remodeling after MI.

### CIP4 gene targeting reduces I/R injury

CaNAβ promotes pathological cardiac remodeling but is also cardioprotective.^9,17^ Therefore, we next considered whether despite CIP4 CKO’s benefit in chronic remodeling, CIP4 CKO would be detrimental in acute MI induced by I/R injury. As measured by 4D echocardiography and corroborated by strain measurements (Figures 2A-D and S4, Videos S7-15), Injury induced by transient left coronary artery ligation for 30 minutes and 24 hours reperfusion resulted in systolic dysfunction with minimal ventricular dilatation (increased end-diastolic volume [EDV]) for the three CIP4 CKO, MCM and *CIP4^f/f^* cohorts, including increased end-systolic volume (ESV) and decreased ejection fraction (EF) . Notably, after I/R injury, EF was 7% and 13% higher for the CIP4 CKO cohort than for the MCM and *CIP4^f/f^* controls, respectively. This correlated with improvement in global endocardial circumferential strain for the CIP4 CKO cohort when compared to the control cohorts (CIP4 CKO – -23.8±1.1% vs. *CIP4^f/f^* - -16.3±0.6%, p < 0.0001; vs. MCM – -21.0±0.9%, *p* = 0.054, Figure S4H).

**Figure 2.**
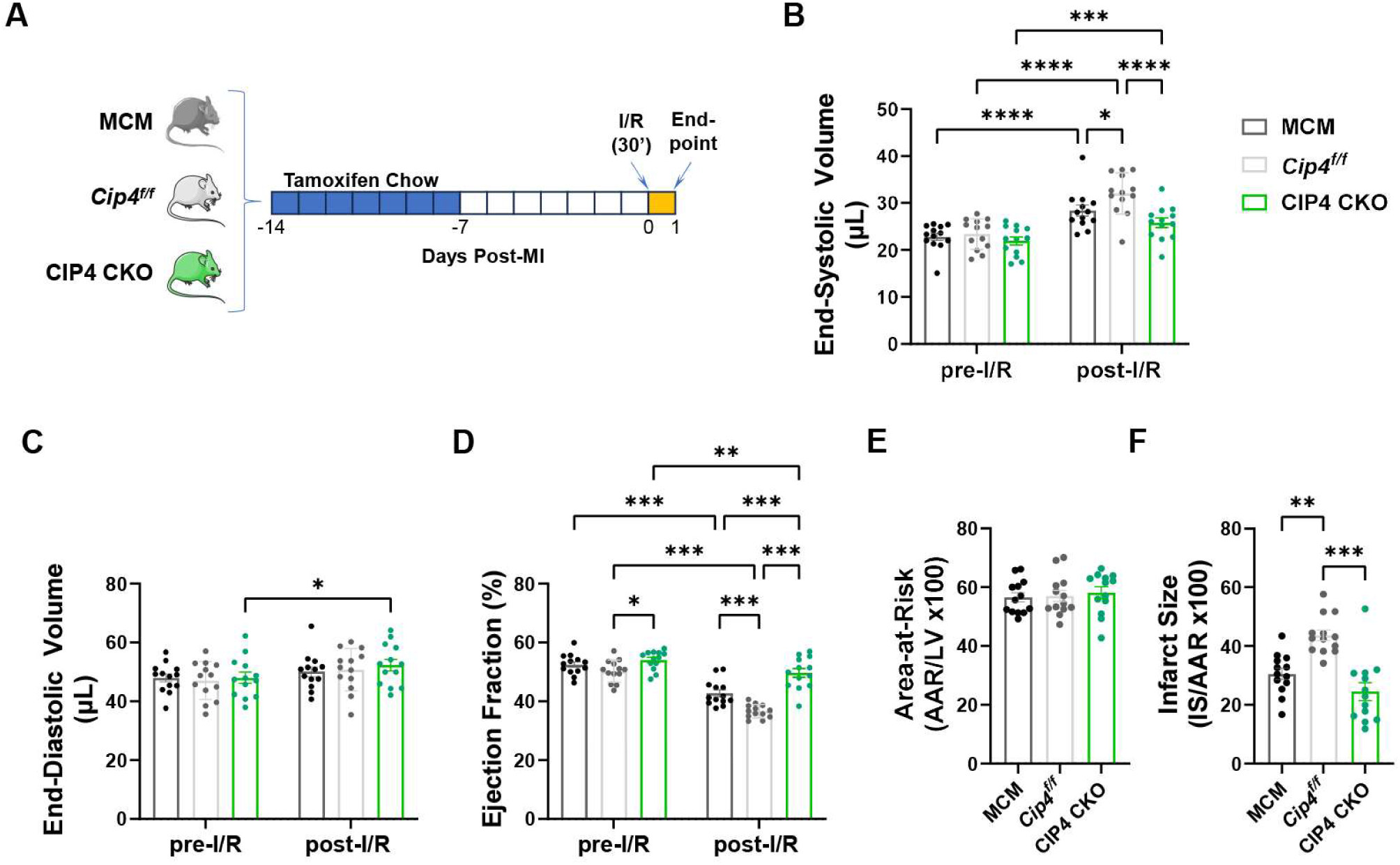
Cardiac function after ischemia/reperfusion injury is improved by *CIP4* conditional gene targeting. **A.** 7-8-week-old male and female CIP4 CKO *CIP4^f/f^*;Tg(Myh6-cre/Esr1*) and control Tg(Myh6-cre/Esr1*) (MCM) and *CIP4^f/f^* mice were treated with tamoxifen-laden chow for one week to induce cre-catalyzed recombination, followed by one week on normal chow before being subjected to transient ligation of the left coronary artery for 30 minutes and 24 hours reperfusion. **B-D.** LV end-systolic volume, end-diastolic volume and ejection fraction by 4D imaging. *n* = 13 for each cohort. Data analyzed by matched 2-way ANOVA and Tukey’s multiple comparison test. **E-F.** Area-at-risk and infarct size by histological Evan’s Blue and 2,3,5-triphenyltetrazolium chloride staining. *n* = 13 for each cohort. Data in E analyzed by 1-way ANOVA and Tukey’s multiple comparison test; data in F analyzed by Kruskal-Wallis and Dunn’s multiple comparison tests. \**p*<0.05; \*\**p*<0.01; *** *p*<0.001; **** *p*<0.0001.

To directly test for a cardioprotective role for CIP4-CaNAβ signalosomes, infarct size was measured by Evan’s Blue and 2,3,5-Triphenyltetrazolium chloride (TTC) histological staining (Figure S2). Importantly, while area-at-risk was similar for all 3 cohorts, infarct size was not increased by CIP4 CKO (Figure 2E,F). In addition, CIP4 CKO showed decreased infarct size compared to *CIP4^f/f^* controls cohort, and a trending decrease relative to the MCM controls when measured histologically or by segmental strain analysis (Figures 2E,F and S4D,I). Taken together, these results suggest that in contrast to CaNAβ knock-out,^17^ CIP4 CKO is not deleterious during I/R injury, and, moreover, is functionally beneficial in both acute and chronic MI.

### Cardiomyocyte-Specific Targeting of CaNAβ2 Worsens I/R Injury

It has been reported that CaNAβ gene knock-out worsened I/R injury,^17^ while transgenic cardiomyocyte-specific CaNAβ1 overexpression did not improve cardiac function in acute MI due to I/R injury despite inhibiting late remodeling after MI.^15,28^ These results suggested that CaNAβ2 is the primary mediator of cardioprotection in I/R injury. However, even though CIP4 associated CaN activity is dependent upon CaNAβ2 expression (and PP-dependent CIP4 anchoring),^16^ as shown above, CIP4 CKO was not deleterious in I/R injury. To test directly whether CaNAβ2 serves any role in cardioprotection, we assessed the effects of cardiomyocyte-specific CaNAβ2 inhibited expression in I/R injury. Serotype 9 adeno-associated virus (AAV9) were generated to express in cardiomyocytes CaNAβ2 (shCaNAβ2) or control (shControl) *MIR30A* small hairpin RNA (shRNA), and AAV9-transduced wildtype C57BL/6N mice were subjected to I/R injury (Figure 3A,B). LV EF was decreased after I/R injury 12% for the shCaNAβ2 cohort compared to 5% for the shControl control (Figures 3C-E and Videos S16-17). The worse systolic dysfunction of the infarcted shCaNAβ2 cohort was corroborated by measurement of LV global and segmental circumferential and longitudinal strain (Figure S5). Histological assay showed that shCaNAβ2 increased infarct size 78% over shControl mice (Figure 3F,G). Measurement of dysfunctional myocardium by segmental analysis of either circumferential and longitudinal strain was consistent with increased injured myocardium in the presence of CaNAβ2 shRNA (Figure 3H,I and S5D). In sum, these results suggest that in contrast to CIP4, CaNAβ2 expression is important for cardioprotection during I/R injury.

**Figure 3.**
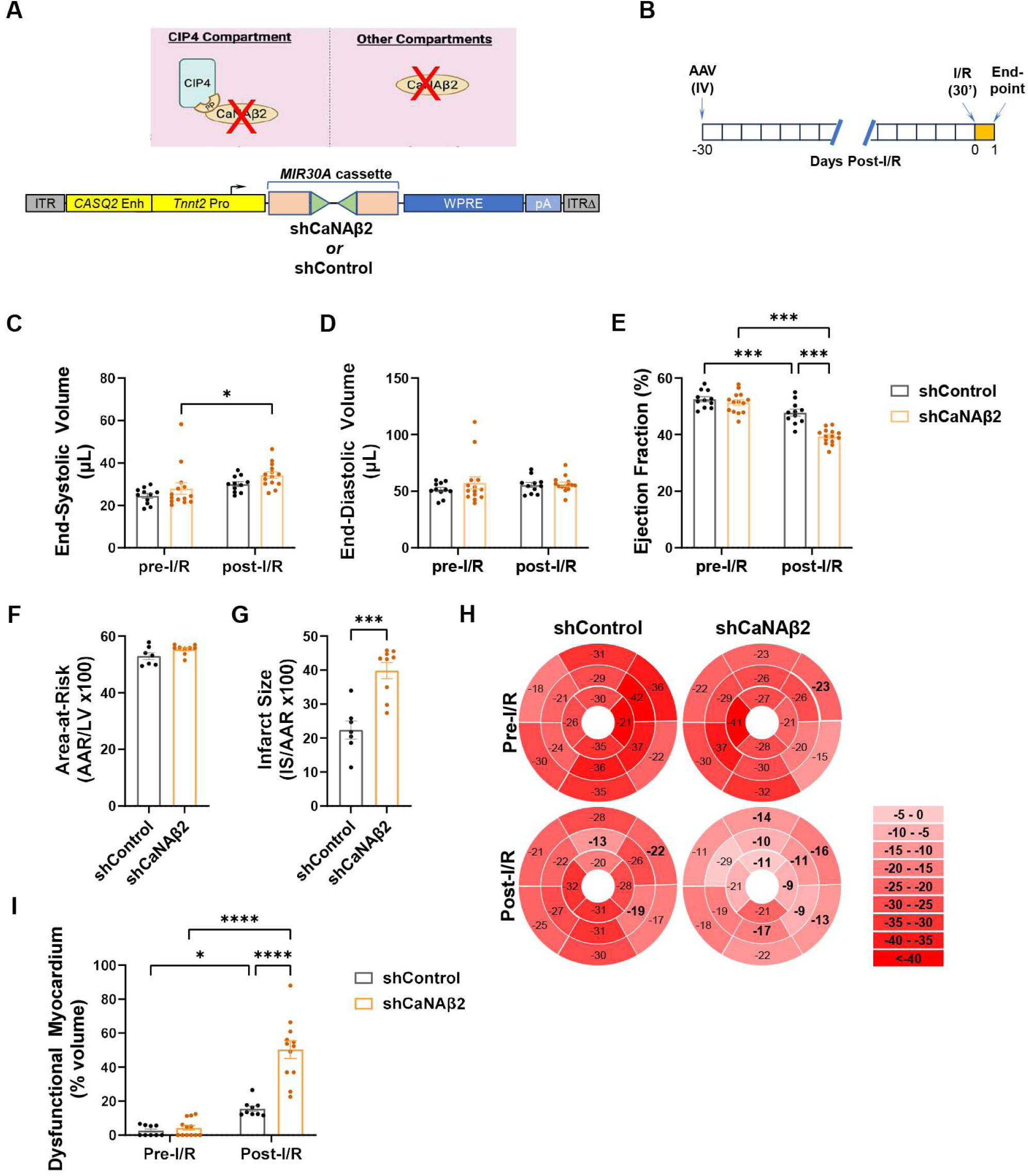
Calcineurin Aβ2 is cardioprotective during ischemia/reperfusion injury. **A.** *top,* CaNAβ2 was depleted throughout the cardiomyocyte by RNA interference. *bottom,* Self-complementary AAV9 vectors were used to express MIR30A control and CaNAβ2 shRNA minigenes^48^ under the direction of a cardiomyocyte-specific chicken cardiac troponin T (*TNNT2*) promoter and a human calsequestrin (*CASQ2*) enhancer.^49,50^ ITR – inverted terminal repeat (Δ – deleted); WPRE - woodchuck hepatitis virus post-transcriptional regulatory element; pA - SV40 polyadenylation signal. **B.** 1-month-old male and female C57BL/6NJ mice were injected with 10^12^ vg AAV (∼7 × 10^13^ vg/kg) 4 weeks before being subjected to transient ligation of the left coronary artery for 30 minutes and 24 hours reperfusion. **C-E.** LV end-systolic volume, end-diastolic volume and ejection fraction by 4D imaging. *n* = 11 (shControl) and 14 (shCaNAβ2). Data analyzed by matched 2-way ANOVA and Uncorrected Fisher’s LSD test. **F,G.** Area-at-risk and infarct size by histological Evan’s Blue and 2,3,5-triphenyltetrazolium chloride staining. *n* = 7 (shControl) and 9 (shCaNAβ2). Data analyzed by two-tailed t tests. **H.** Representative bullseye plots for circumferential peak systolic strain analysis of PS-SAX images. Sectors labelled with bolded text indicate regions with decreased strain. **I.** Dysfunctional myocardium as assessed by circumferential peak systolic strain analysis and expressed as % LV volume. *n* = 9 (shControl) and 12 (shCaNAβ2). Data analyzed by matched 2-way ANOVA and Uncorrected Fisher’s LSD test. \**p*<0.05; *** *p*<0.001; **** *p*<0.0001.

### Expression of VIVIT and CaNAβ PP CaN anchoring disruptor peptides differentially affect outcome after ischemia/reperfusion injury

Previously published live cell imaging results suggest that CIP4-bound CaNAβ2 comprises a minority of CaNAβ in the cardiomyocyte.^16^ We, therefore, considered that CIP4-bound CaNAβ2 might be unrelated to the cardioprotective action of CaNAβ2 located elsewhere in the myocyte. Expression of a polyproline (PP) peptide based upon the unique N-terminus of CaNAβ will compete the binding of CaNAβ2 to CIP4, and, like CIP4 CKO, expression of a PP-GFP fusion protein improved pathological cardiac remodeling in response to chronic pressure overload.^16^ To test whether CIP4-bound CaNAβ2 contributes to cardioprotection, adult wildtype C57BL/6N mice were administered AAV9 vectors that express in cardiomyocytes PP-GFP or GFP control (Figures 4A and S6A). The mice were studied 1 month later by I/R injury (Figure 4B). For comparison, a VIVIT-GFP fusion protein was also expressed using AAV9. The VIVIT peptide is based upon a PxIxIT short linear motif present within scaffold proteins and substrates like NFAT that bind a conserved allosteric docking site on all CaN catalytic A-subunits.^7^ Expression of the VIVIT peptide has been shown to inhibit NFAT activation and myocyte survival *in vitro*.^17,23,24^

**Figure 4.**
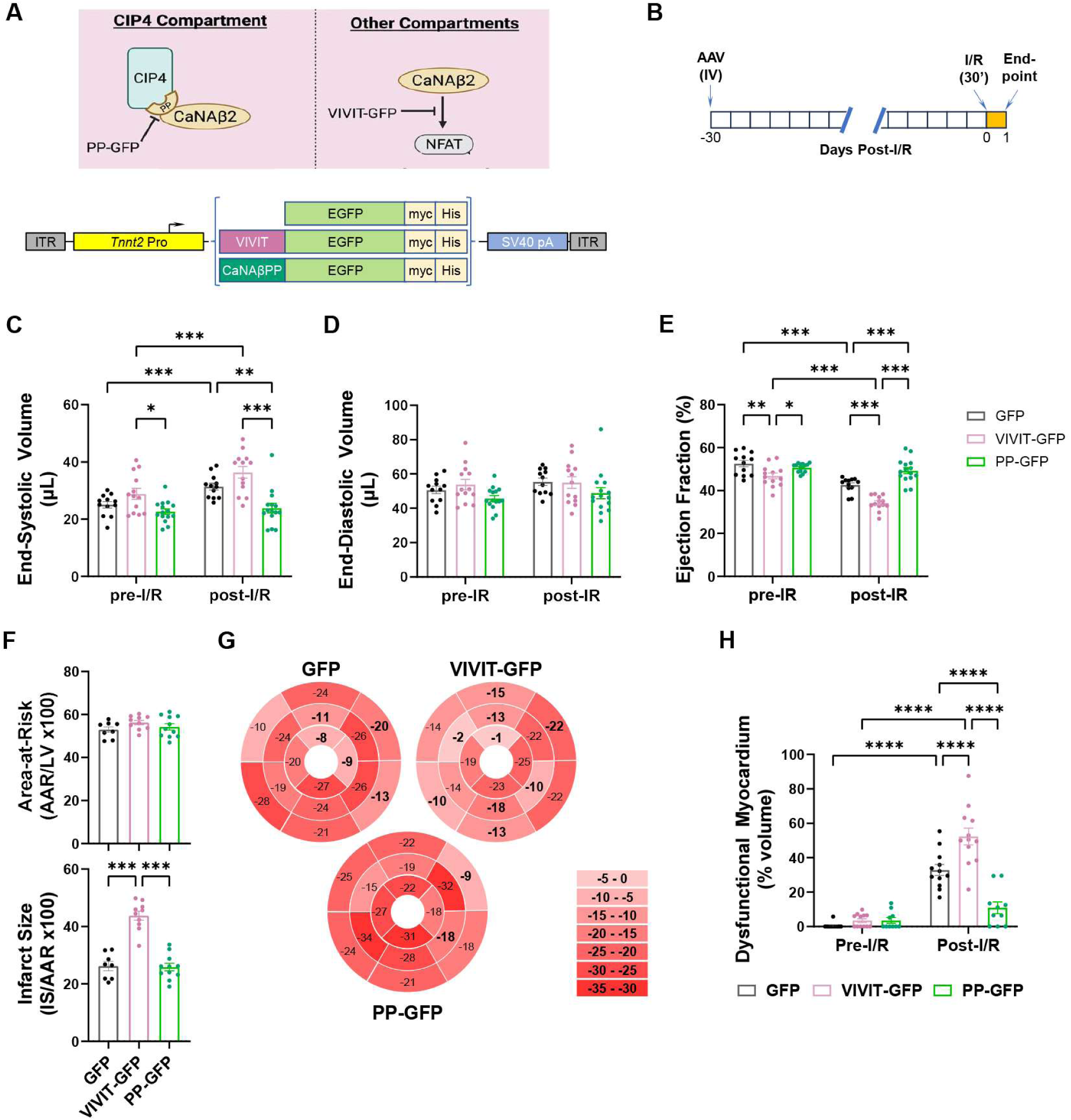
CaNAβ PP and VIVIT calcineurin anchoring disruptor peptides improve and worsen outcome after ischemia/reperfusion injury, respectively. **A.** *top,* PP-GFP blocks CIP4-CaNAβ binding, while VIVIT-GFP inhibits CaN-NFAT signaling. CIP4-CaNAβ signaling is not associated with NFAT activation.^16^ *bottom,* AAV9 vectors were used to express VIVIT-GFP, PP-GFP and GFP control fusion proteins under the direction of a cardiomyocyte-specific chicken cardiac troponin T (*TNNT2*) promoter.^50^ ITR – inverted terminal repeat; SV40 pA - SV40 polyadenylation signal. **B.** 1-month-old male and female C57BL/6NJ mice were injected with 10^12^ vg AAV (∼7 × 10^13^ vg/kg) 4 weeks before being subjected to transient ligation of the left coronary artery for 30 minutes and 24 hours reperfusion. **C-E.** LV end-systolic volume, end-diastolic volume and ejection fraction by 4D imaging. *n*: GFP – 12; VIVIT-GFP – 13; PP-GFP - 15. Data analyzed by matched 2-way ANOVA and Tukey’s multiple comparison test. **F.** Area-at-risk and infarct size by histological Evan’s Blue and 2,3,5-triphenyltetrazolium chloride staining. *n*: GFP – 8; VIVIT-GFP – 10; PP-GFP - 11. Data analyzed by Data analyzed by 1-way ANOVA and Tukey’s multiple comparison test. **G.** Representative bullseye plots for circumferential peak systolic strain analysis of PS-SAX images. Sectors labelled with bolded text indicate regions with decreased strain. **H.** Dysfunctional myocardium as assessed by circumferential peak systolic strain analysis and expressed as % LV volume. *n*: GFP – 12; VIVIT-GFP – 12; PP-GFP - 10. Data analyzed by matched 2-way ANOVA and Tukey’s multiple comparison test. \**p*<0.05; \*\**p*<0.01; *** *p*<0.001; **** *p*<0.0001.

Compared to GFP, VIVIT-GFP expression induced mild systolic dysfunction (EF: 47±1% vs. 53±2%, *p* = 0.003) 1 month after AAV9 administration (Figures 4C-E), while PP-GFP expression had no significant effect in uninjured mice. Following I/R injury, VIVIT-GFP expression resulted in a 13% further decrease in EF, while EF decreased 10% for control GFP-treated mice (Videos S18-20). In contrast, PP-GFP expression preserved both ESV and EF. The worsening and preservation of systolic function by VIVIT-GFP and PP-GFP expression, respectively, was corroborated by measurement of global and segmental circumferential and longitudinal peak strain (Figure S6). VIVIT-GFP worsened infarct size measured histologically 67%, like shCaNAβ2, while infarct size was not different for PP-GFP- and GFP-treated mice (Figure 4F). In contrast, the fraction of dysfunctional myocardium measured by segmental circumferential and longitudinal peak strain analysis was dramatically less following I/R injury for the PP-GFP cohort, while, consistent with histological measurements, increased by VIVIT-GFP expression (Figures 4G-H and S6E). Taken together, these results suggest that VIVIT-GFP exacerbated I/R injury, while PP-GFP preserved cardiac function in the absence of histologically detectable cardioprotection.

### Inhibition of CaNAβ PP anchoring improves cardiac structure and function in chronic ischemic cardiomyopathy

As PP-GFP expression and CIP4 CKO improved systolic function after I/R injury, we next tested whether like CIP4 CKO, PP-GFP might also be beneficial in chronic ischemic cardiomyopathy. Wildtype male C57BL/6N mice were subjected to LCA PL or sham operation and randomized 2 days later by EF for treatment with AAV9.PP-GFP, AAV9.VIVIT-GFP, or AAV9.GFP control (Figure 5A,B). At endpoint 8 weeks after survival surgery, the sham-operated cohorts were similar in cardiac structure and function (Figure 5C-E, Videos S21-23). During the 8 weeks post-MI, the GFP-treated infarcted cohort showed a steady decline in systolic function, evident as a 6% drop in EF (*p* = 0.004, Video S24). The VIVIT-GFP treated MI cohort exhibited a more profound phenotype, with a 14% drop in EF and a 47 µL and 48 µL increase in ESV and EDV, respectively (*p* < 0.0001 for each parameter, Video S25). In contrast, the PP-GFP treated MI cohort exhibited no LV dilatation and, notably, an 11% increase in EF (*p* < 0.0001) to 44±4% at endpoint that correlated with improvements in stroke volume and cardiac output (Figure S7A-E, Video S26). Segmental and global circumferential and longitudinal strain analysis confirmed these differences in endpoint systolic function (Figure S7F-L).

**Figure 5.**
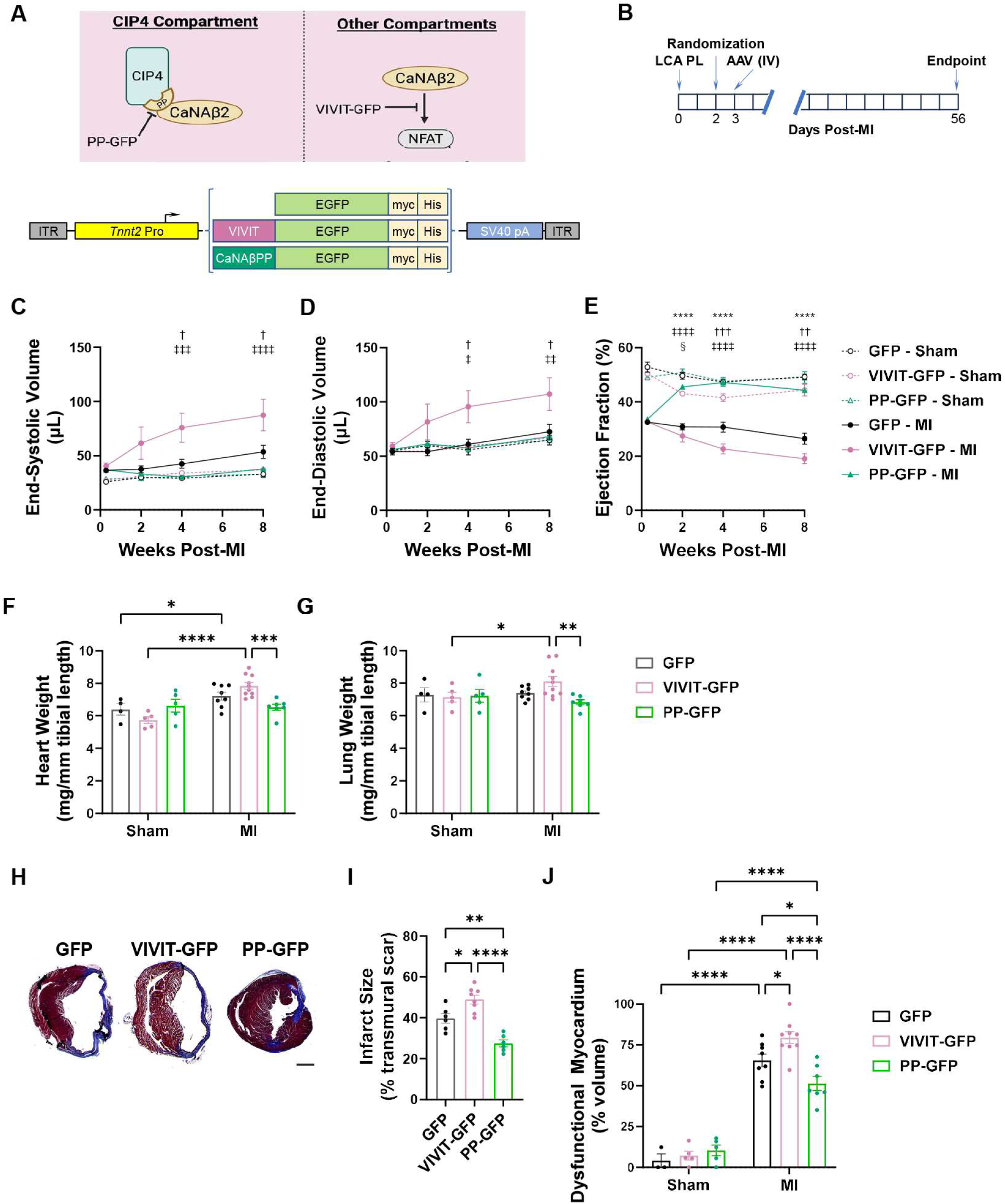
CaNAβ PP and VIVIT calcineurin anchoring disruptor peptides improve and worsen outcome in chronic ischemic cardiomyopathy, respectively. **A.** *top,* PP-GFP blocks CIP4-CaNAβ binding, while VIVIT-GFP inhibits CaN-NFAT signaling. CIP4-CaNAβ signaling is not associated with NFAT activation.^16^ *bottom,* AAV9 vectors were used to express VIVIT-GFP, PP-GFP and GFP control fusion proteins under the direction of a cardiomyocyte-specific chicken cardiac troponin T (*TNNT2*) promoter.^50^ ITR – inverted terminal repeat; SV40 pA - SV40 polyadenylation signal. **B.** 9-11-week-old male C57BL/6NJ mice were subjected to permanent ligation of the left coronary artery (LAD PL), randomized 2 days later by ejection fraction acquired by 4D echocardiography, and the following day injected with 10^12^ vg AAV (∼4 × 10^13^ vg/kg). **C-E.** LV end-systolic volume, end-diastolic volume and ejection fraction by serial 4D echocardiography. *n*: GFP Sham – 4; VIVIT-GFP Sham – 5; PP-GFP Sham – 4; GFP MI – 7; VIVIT-GFP MI – 9; PP-GFP MI - 7. Data analyzed by matched 2-way ANOVA (mixed-effects model) and Tukey’s multiple comparison test. * *p*-values for GFP MI vs. PP-GFP MI; ^†^ *p*-values for GFP MI vs. VIVIT-GFP MI; ^‡^ *p*-values for PP-GFP MI vs. VIVIT-GFP MI; **^§^** *p*-values for GFP Sham vs. VIVIT-GFP Sham. **F,G.** Heart and wet lung weight indexed to tibial length by gravimetric measure. *n*: GFP Sham – 4; VIVIT-GFP Sham – 5; PP-GFP Sham – 5; GFP MI – 8; VIVIT-GFP MI – 10; PP-GFP MI - 7. Data analyzed by Data analyzed by 2-way ANOVA and Tukey’s multiple comparison test. **H.** Representative transverse cardiac sections stained with Masson’s Trichrome. Scale bar – 1 mm. **I.** Infarct size measured as % left ventricular circumference containing >50% transmural scar. *n*: GFP – 6; VIVIT-GFP – 8; PP-GFP MI - 6. Data analyzed by 1-way ANOVA and Tukey’s multiple comparison test. **J.** Dysfunctional myocardium as assessed by circumferential peak systolic strain analysis and expressed as % LV volume. *n*: GFP Sham – 3; VIVIT-GFP Sham – 5; PP-GFP Sham – 4; GFP MI – 8; VIVIT-GFP MI – 9; PP-GFP MI - 7. Data analyzed by 2-way ANOVA and Tukey’s multiple comparison test. Repeated symbols indicate: \**p*<0.05; \*\**p*<0.01; *** *p*<0.001; **** *p*<0.0001.

Histological assessment of infarct size showed that VIVIT-GFP increased 23%, while PP-GFP decreased 31% infarct size when compared to GFP control (Figure 5H,I). Measurement of fractional dysfunctional myocardium by segmental circumferential and longitudinal strain analysis corroborated these results (Figures 5J and S7I). In sum, expression of VIVIT-GFP, which inhibits CaN-NFAT signaling, worsened cardiac function and structure after MI due to LCA PL. In contrast, PP-GFP anchoring disruption, which inhibits the formation of CIP4-CaNAβ2 signalosomes, markedly improved cardiac function and structure in chronic ischemic cardiomyopathy. These results support the hypothesis that in contrast to the broad-based targeting of CaNAβ2, selective inhibition of CIP4-CaNAβ2 signalosomes is beneficial in ischemic heart disease.

### Lack of effect by CaNAβ PP anchoring disruption on T-cells in vitro

CaNAβ gene targeting has been reported to inhibit T-cell development and function in young mice, overlapping in phenotype with cyclosporin A-induced immunosuppression.^18^ In order to screen for a possible immunosuppressive effect of PP-anchored CaNAβ in T-cells as a potential off-target effect of its cardioprotective strategy, we transduced primary mouse CD4^+^ T-cells, which express CIP4,^29^ with lentivirus that expresses PP-GFP or GFP control. As the degree of T-cell receptor stimulation dictates T-cell activation outcomes and T-cells are central to cardiac repair and ischemic heart failure,^30^ T-cells were stimulated by high and low doses of αCD3 and αCD28 activating antibodies. T-cells were transduced with similar efficacy as shown by flow cytometry for GFP expression (40±17 and 36±17 % [mean ± SD] of the GFP and PP-GFP lentivirus-transduced cells treated with the high dose of activating antibodies, respectively, Figure S8). Viability was also not different between PP-GFP and GFP transduced T-cells (Figure 6B). Importantly, surface expression of early (CD69) and late (CD25) markers of T-cell activation were increased in GFP-expressing cells in an αCD3 and αCD28 dose-dependent manner, regardless of expression of the PP anchoring disruptor peptide (Figure 6C-E). Likewise, mRNA levels of cytokines resulting from T-cell activation, such as IL-2, IFNγ, IL-17, and TNFα, were similarly elevated in both groups regardless of GFP or PP-GFP expression (Figure 6F). These data demonstrate that CaNAβ PP anchoring does not serve a significant role in T-cell activation and, moreover, suggest that a heart failure drug targeting CIP4-CaNAβ signalosomes would not be associated with immunosuppression.

**Figure 6.**
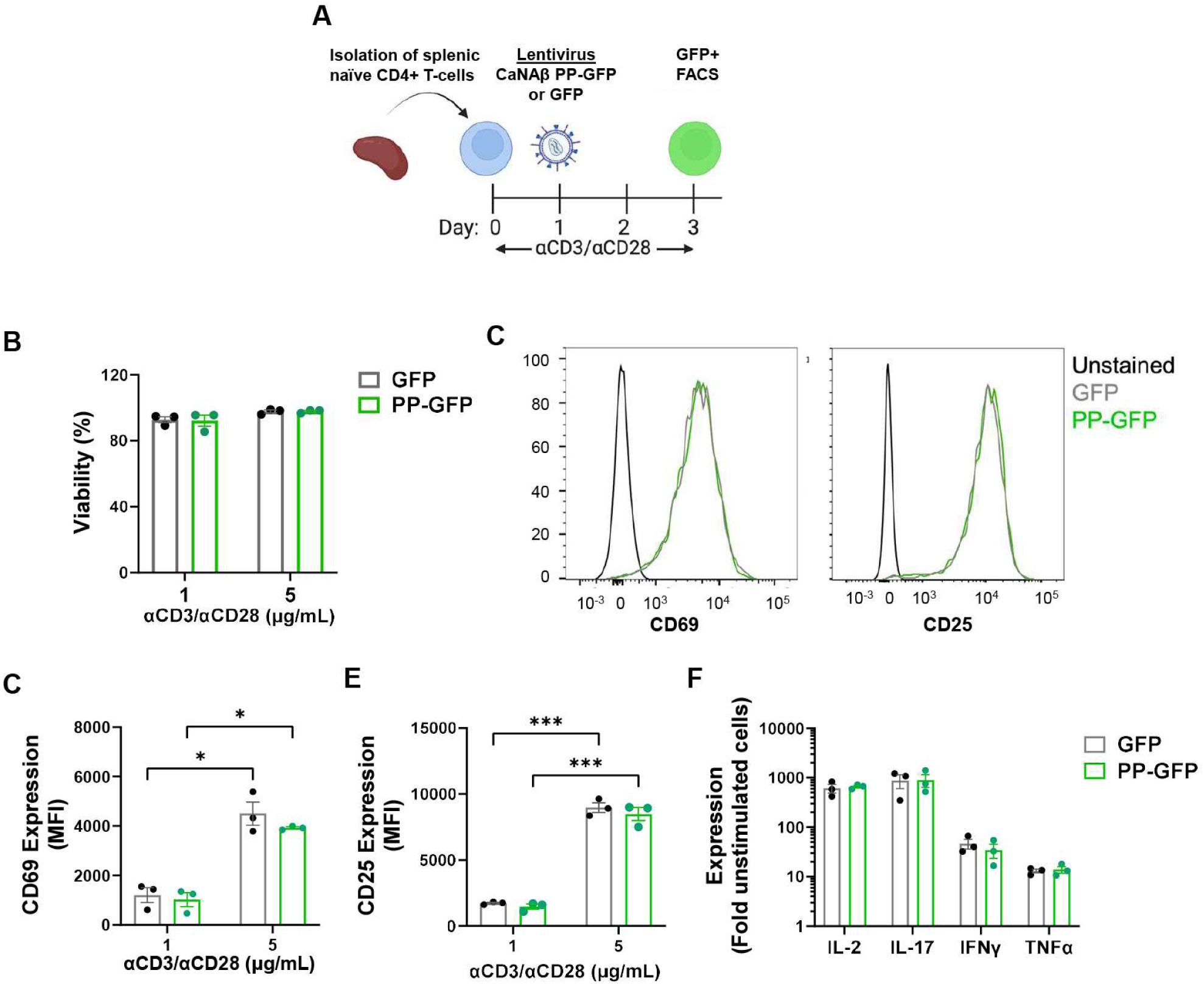
PP anchoring disruption does not affect T-cell activation. **A.** Naïve CD4+ T-cells were isolated from wild type C57B6/J mice and stimulated with 1 or 5 µg/mL αCD3 and αCD28 before transduction with lentivirus that express PP-GFP or GFP control and fluorescence-activated cell sorting (FACS) analysis and RNA isolation. **B.** Cell viability by 7-amino-actinomycin D (7AAD) stain. **C.** Representative FACS analysis for CD69 and CD25 mean fluorescence intensity (MFI) for 5 µg/mL αCD3/CD28-stimulated group. **D,E.** Quantification of CD69 and CD25 MFI. CD69 and CD25 MFI for unstimulated, naïve cells were 66±11 and 124±14 (mean±SEM, not shown), respectively. **F.** qRT-PCR for fold gene expression relative to naïve cells. Data analyzed by matched 2-way ANOVA and Uncorrected Fisher’s LSD tests. *n* = 3 independent T-cell preparations. \**p*<0.05; *** *p*<0.001.

## Discussion

CaNAβ has been established by gene knock-out as an important mediator of both pathological cardiac hypertrophy and cardiomyocyte survival, as well as serving important roles in the immune system and other tissues.^5,6,31^ This pleiotropy has been a roadblock to the development of CaN-targeted therapeutics for heart failure, particularly in the context of chronic ischemic cardiomyopathy where this a significant risk of recurrent myocardial infarction.^1^ We have shown that the endosomal scaffold protein CIP4 binds CaNAβ via its PP-domain, constituting an independent Ca^2+^ and CaNAβ2 signaling compartment that promotes myocyte hypertrophy and pathological cardiac remodeling in response to chronic pressure overload.^16^ The results presented herein demonstrate that in contrast to generalized CaNAβ2 depletion or inhibition of CaN interactions with the large family of PxIxIT motif-containing substrates that include NFAT transcription factors,^7^ CIP4 gene targeting and PP-anchoring disruption are not deleterious and, instead, can be functionally beneficial during I/R injury (Figure 7). Notably, both cardiomyocyte-specific CIP4 knock-out and PP-GFP expression improved cardiac structure and function in chronic ischemic cardiomyopathy. As *in vitro* results suggest that PP-directed CaNAβ signaling is irrelevant to T-cell function, CIP4 signalosomes may constitute a favorable target for intervention in both acute MI and chronic pathological cardiac remodeling under diverse pathophysiological conditions.

**Figure 7.**
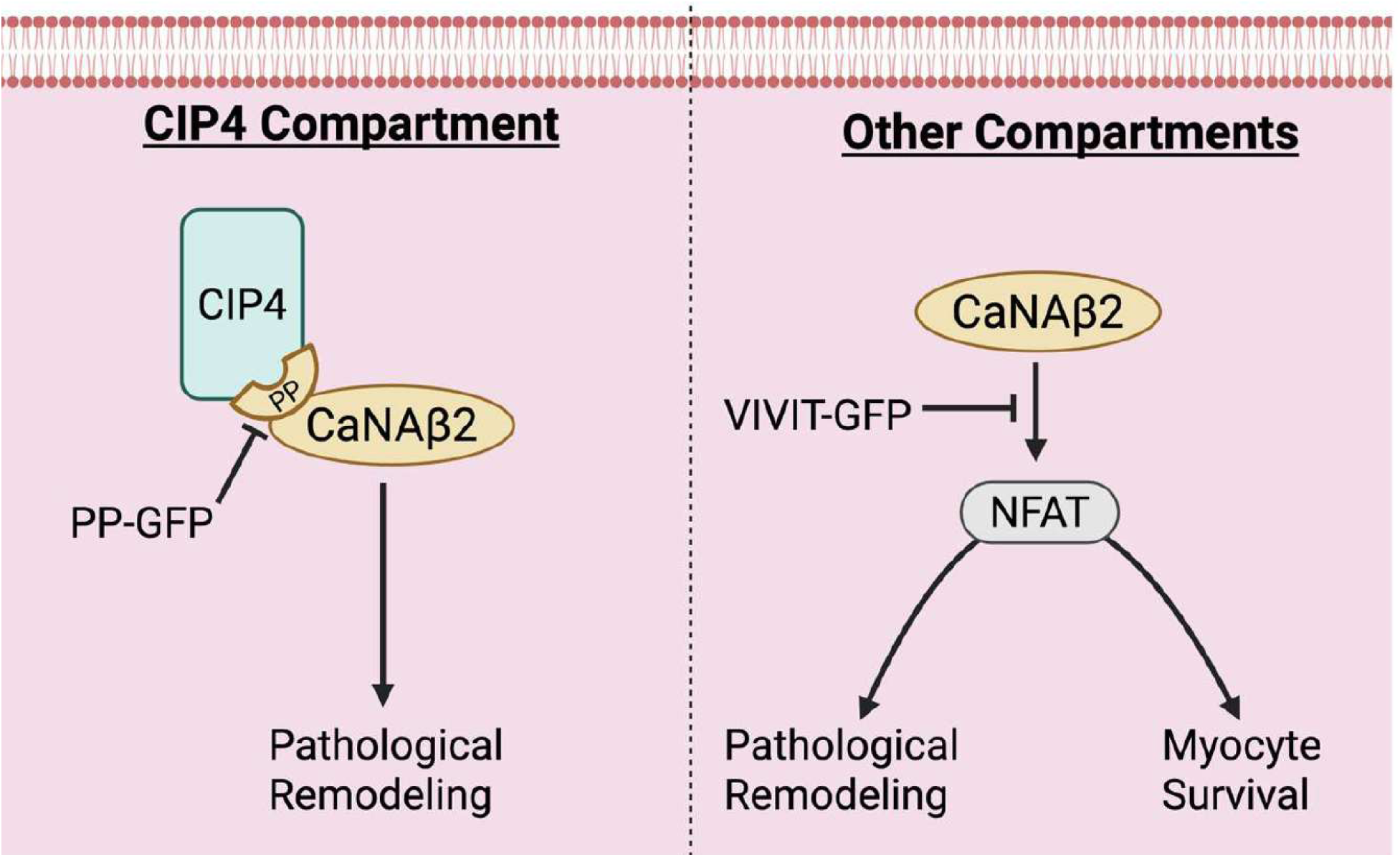
Model for compartmentalized CaNAβ2 signaling in cardiac myocytes. CIP4 organizes a CaNAβ2 compartment that promotes pathological cardiac remodeling in both ischemic and pressure overload disease.^16^ CaNAβ2 anchored by its PP domain to CIP4 can be targeted via expression of a PP-GFP anchoring disruptor fusion protein. CaNAβ2 in other myocyte intracellular compartments promotes both pathological cardiac remodeling and myocyte survival (cardioprotection) via NFAT activation.^9,16,17^ Expression of VIVIT-GFP inhibits CaN activation of NFAT,^24^ while PP-anchored CaNAβ is not associated with NFAT activation.^16^ Results obtained using CaNAβ2 shRNA, which will inhibit CaNAβ2 levels throughout the cell, but phenocopies the VIVIT-GFP peptide, indicate that loss of CaNAβ2 cardioprotection is dominant to any beneficial effects in ischemic disease of inhibiting CaNAβ2 function in pathological remodeling.

In this project, we took advantage of recent advances in 4D echocardiography and strain analysis to corroborate histological results following I/R injury and chronic MI. MI due to I/R injury is traditionally measured by Evan’s Blue and TTC histological staining, which relies on colorimetric differences to quantify perfused, viable, and non-viable myocardium.^32^ Chronic MI can be quantified in Masson’s Trichrome stained fixed tissue by the measurement of blue scar tissue. As infarcted tissue is hypocontractile, MI can also be identified as regions exhibiting reduced longitudinal or circumferential strain using high-resolution echocardiography.^33,34^ Dysfunctional myocardium measured as % endocardial length in PS-LAX views or % volume using SAX images correlated with histological measurement of infarct size both in acute and chronic MI, with the notable exception of the experiment involving PP-GFP expression and I/R injury. While VIVIT-GFP expression resulted in worse I/R injury than either PP-GFP or GFP regardless of quantification method, PP-GFP resulted in no difference in I/R injury when measured histologically but significantly improved I/R injury when measured by strain. It is possible that the different results reflect a relative lack of sensitivity in the histological measurement of infarction post-I/R. Alternatively, the strain measurements used in this study are not designed to distinguish stunned and non-viable myocardium.^35^ It is possible that the improvement in systolic function following treatment with AAV9.PP-GFP, whether measured by ESV, EF, or strain, reflects decreased stunning relative to the VIVIT-GFP and GFP cohorts as opposed to improved myocardial survival. Parenthetically, we note that despite the similarity in phenotype of MCM and *CIP4^f/f^* controls in uninjured and sham-operated mice, as well as in chronic MI following LCA PL, MCM control mice had decreased infarct size and improved cardiac function after I/R injury when compared to *CIP4^f/f^* control littermates (Figures 1 and 2). Tamoxifen-treated MCM mice have been reported to exhibit transient systolic dysfunction, which is absent at the low dose of tamoxifen used here.^36^ In contrast, the present results suggest that a low dose of tamoxifen in conjunction with the MCM transgene may induce a preconditioning-like phenomenon, underscoring the need for appropriate controls in genetic studies.

It is well-established that CaN phosphatase activity is important for cardiac myocyte survival and hypertrophy via activation of NFAT transcription factors.^6^ Early on, expression of a constitutive active truncated CaNAα was shown to decrease cardiomyocyte cell death following I/R injury in mice.^37^ This improvement in cardiac survival was in contrast to its induction of pathological cardiac hypertrophy and heart failure in uninjured transgenic mice.^38^ Conversely, via decreased NFAT activation, infarct size and myocyte apoptosis following I/R injury were found to be increased in global, constitutive CaNAβ knock-out mice,^17^ which also exhibited decreased pressure overload-induced pathological hypertrophy.^9^ Notably, transgenic expression of the minor CaNAβ isoform CaNAβ1, that is localized by palmitoylation to the plasma membrane and Golgi by its unique C-terminus,^39^ improved cardiac function and reduced scar size after MI induced by permanent coronary artery ligation.^15^ CaNAβ1, however, activated AKT, ATF4, and serine and one-carbon metabolism, but not NFAT, and was anti-hypertrophic.^15,40^ We show here that like CaN isoform-non-specific VIVIT-mediated inhibition, depletion by RNA interference of CaNAβ2 exacerbated I/R injury in a myocyte autonomous manner (Figures 3 and 4). Taken together, these results support a role for CaN in cardioprotection, and, moreover, suggest that loss of cardiomyocyte CaNAβ2 is a major contributor to the worsened phenotype in I/R injury conferred by global CaNAβ gene targeting.

Paradoxically, CaN signaling can also be pro-apoptotic.^41^ By binding the CaNAβ PP domain, CIP4 binds mainly, if not exclusively, CaNAβ2 in the adult cardiomyocyte.^16^ Consistent with the association of CaN with diverse scaffold proteins,^5,7^ results previously obtained by live cell imaging suggest that CIP4-associated CaNAβ2 comprises a small, localized pool of CaN within the myocyte.^16^ Although CIP4-CaNAβ2 signalosomes are pro-hypertrophic, CIP4-CaNAβ2 activity has not been associated with NFAT activation.^16^ Neither PP-GFP nor CIP4 CKO was found to increase infarct size after I/R injury. These results contrast with those for VIVIT-GFP in I/R injury, which inhibits NFAT activation.^24^ Instead, both PP-GFP and CIP4 CKO improved cardiac structure and function long term after MI. While we have defined the physiological relevance of CIP4-CaNAβ2 signalosomes, how CIP4-bound CaNAβ2 promotes adverse remodeling remains unclear. Future research will focus on identifying which CaN substrates contribute to adverse CIP4 signaling in that subcellular compartment.

The results of CaNAβ and CIP4 knock-out and PP-GFP expression in chronic disease models, including the results shown here for chronic ischemic cardiomyopathy, support a role for CaNAβ2 in pathological cardiac remodeling,^9,16^ whether through CIP4 or in other subcellular compartments (Figure 7). In contrast to the beneficial effects of VIVIT peptide in pressure overload hypertrophy,^25^ the worse outcome following VIVIT-GFP in the chronic MI model presumably reflects the unique importance of CaNAβ2-mediated cardioprotection in ischemic disease. Likewise, the exacerbation of I/R injury by shCaNAβ2 stands in stark contrast to the lack of increased infarct size and the improved cardiac function for CIP4 CKO and PP-GFP-expressing mice in the I/R model. A limitation of this study is that PP-GFP expression was used to probe the function of CIP4-associated CaNAβ. PP-GFP would be expected to compete CaNAβ binding to other scaffolds which bind the PP domain, as well the binding to CIP4 of other CIP4 SH3 domain binding partners.^16^ While we hypothesize that CIP4-bound CaNAβ2 does not contribute to cardioprotection like CaNAβ2 in other intracellular compartments, it is also possible that CaNAβ2 is simply not associated with CIP4 during I/R and that PP-GFP blockade of binding of other protein partners to the CIP4 SH3 domain confers the beneficial phenotype in that model. Regardless, that neither CIP4 CKO nor PP-GFP promoted myocardial loss after I/R injury supports the safety of targeting CIP4-CaNAβ2 signalosomes complexes in pathological cardiac remodeling and chronic cardiac disease.

Current therapy for acute MI is generally focused on coronary reperfusion,^42^ and targeting of CIP4-CaNAβ2 signalosomes, which might enhance myocardial function, could be advantageous in this setting. In addition, together with earlier findings in the pressure overload model,^16^ results obtained followed permanent coronary artery ligation constitute proof-of-concept that targeting of CIP4-CaNAβ2 signalosomes can be efficacious for the prevention of heart failure arising from diverse etiologies. A cardiac specific AAV gene therapy could be effective for the prevention or treatment of heart failure, but due to the latency of AAV therapies, a small molecule inhibiting the CIP4-CaNAβ protein-protein interaction would presumably be preferable for the treatment of acute MI. This approach would represent a distinct therapeutic strategy compared to currently available CaN inhibitors such as the immunosuppressants cyclosporin A and FK506 (tacrolimus). While instrumental for the wide-spread deployment of solid organ transplant therapies,^31^ CaN inhibitors currently in use clinically have a variety of off-target effects, including prominent renal toxicity.^43^ Interestingly, CaN inhibitor-associated nephrotoxicity has been associated with inhibition of renal CaNAα.^44^ CaNAβ is also important for T-cell development and function, such that CaNAβ knock-out mice have defective allograft rejection.^18^ Young CaNAβ knock-out mice have reduced CD3+ T-cells, as well as CD4+ and CD8+ cells.^18,45^ In addition, T-cells isolated from CaNAβ knock-out mice have reduced proliferation in response to αCD3 and αCD28 and inhibited IL-2 expression in response to phorbol ester and ionomycin.^10,18^ Our results demonstrate that *in vitro* T cell activation and viability are similar regardless of PP-anchored CaNAβ. While further studies will be necessary regarding potential off-target effects of PP anchoring disruption therapies, the lack of effect of PP-GFP on T-cell activation suggest that targeting PP-anchored CaNAβ will not be immunosuppressive. This would be necessary for therapeutic translation, as T-cells play a critical role in the acute response to cardiac ischemia.

Like PP-anchored CaNAβ, CIP4 may itself be an acceptable drug target. We have found that CIP4 cardiomyocyte-specific knock-out in the healthy adult mouse induces no obvious phenotype.^16^ In addition, CIP4 global knock-out mice exhibit no obvious anatomical, behavioral, growth, or reproductive pathology, with the exception of a mild thrombocytopenia.^46,47^ Interestingly, CIP4 targeting is potentially beneficial for the treatment of diabetes, and decreased post-prandial serum glucose levels and increased adipocyte glucose uptake and insulin sensitivity were observed for CIP4 knock-out mice, apparently due to decreased glucose transporter 4 internalization.^46^ CIP4 knock-out mice also exhibit normal T- and B-cell development and generally normal immune responses, with defects specifically in T-cell dependent IgG and IgE antibody responses and contact hypersensitivity.^29^ Given the improvement in cardiac structure and function in ischemic and pressure overload cardiomyopathy conferred by CIP4-CaNAβ targeting in mice,^16^ further research is warranted into the translational potential of targeting CIP4-CaNAβ signalosomes.

## Non-standard Abbreviations and Acronyms

AAV: adeno-associated virus
CaN: calcineurin
CaNAβ2: β2 isoform of the calcineurin A-subunit
CIP4: Cdc42-interacting protein 4
CIP4 CKO: CIP4 conditional knock-out
EDV: end-diastolic volume
EF: ejection fraction
ESV: end-systolic volume
F-BAR: Fes-CIP4 homology – Bin/Amphiphysin/Rvs domain
GCS: global circumferential strain
GFP: green fluorescent protein
GLS: global longitudinal strain
I/R: ischemia reperfusion
IFNγ: interferon γ
IL-2: interleukin 2
IL-17: interleukin 17
LCA PL: left coronary artery permanent ligation
LV: left ventricular
MCM: MerCreMer transgene
MI: myocardial infarction
NFAT: nuclear factor of activated T-cells
PE: phenylephrine
PP: CaNAβ polyproline domain
PS-LAX: parasternal long axis
PS-SAX: parasternal short axis
SH3: Src Homology 3
shCaNAβ2: CaNAβ2 small hairpin RNA
scControl: control small hairpin RNA
TNFα: Tumor Necrosis Factor α
TTC: 2,3,5-Triphenyltetrazolium chloride

## Acknowledgments

Drs. Samuelsson and Y. Li performed in vivo experimentation. Drs. Bayer and J. Li and Mr. Lewis performed *in vitro* experimentation. Drs. Kapiloff and Alcaide supervised the research. Dr. Kapiloff conceived the project and wrote the manuscript with the assistance of Drs. Dodge-Kafka and Turcotte and the other co-authors.

## Sources of Funding

This work was supported by NIH Grants R01 HL158052 (Dr. Kapiloff), R01 HL166547 (Dr. Dodge-Kafka), R01 HL144477 and R01 HL165725 (Dr. Alcaide), and F30 HL162200 (Dr. Bayer), and the NHLBI Gene Therapy Resource Program.

## Disclosures

The authors declare no competing financial interests.

## Supplemental Material

Supplemental Methods

Major Resources Table

Tables S1-3

Figures S1-S8

References #51-53.

Video S1-26.

## Notes

### Competing Interest Statement

The authors have declared no competing interest.

